# Adaptive Somatic Mutations Calls with Deep Learning and Semi-Simulated Data

**DOI:** 10.1101/079087

**Authors:** Remi Torracinta, Laurent Mesnard, Susan Levine, Rita Shaknovich, Maureen Hanson, Susan Levine

## Abstract

A number of approaches have been developed to call somatic variation in high-throughput sequencing data. Here, we present an adaptive approach to calling somatic variations. Our approach trains a deep feed-forward neural network with semi-simulated data. Semi-simulated datasets are constructed by planting somatic mutations in real datasets where no mutations are expected. Using semi-simulated data makes it possible to train the models with millions of training examples, a usual requirement for successfully training deep learning models. We initially focus on calling variations in RNA-Seq data. We derive semi-simulated datasets from real RNA-Seq data, which offer a good representation of the data the models will be applied to. We test the models on independent semi-simulated data as well as pure simulations. On independent semi-simulated data, models achieve an AUC of 0.973. When tested on semi-simulated exome DNA datasets, we find that the models trained on RNA-Seq data remain predictive (sens 0.4 & spec 0.9 at cutoff of *P* > = 0.9), albeit with lower overall performance (AUC=0.737). Interestingly, while the models generalize across assay, training on RNA-Seq data lowers the confidence for a group of mutations. Haloplex exome specific training was also performed, demonstrating that the approach can produce probabilistic models tuned for specific assays and protocols. We found that the method adapts to the characteristics of experimental protocol. We further illustrate these points by training a model for a trio somatic experimental design when germline DNA of both parents is available in addition to data about the individual. These models are distributed with Goby (http://goby.campagnelab.org).

## INTRODUCTION

The analysis of high-throughput sequencing data often involves ranking and filtering millions of observed genomic sites to locate candidates of biological or clinical interest. For instance, many approaches and associated software tools have been developed to identify sites of somatic variations in the data derived from the genome of tumors. Methods have been developed for matched DNA samples, where germline and somatic tumor tissues are both available for an individual Pabinger et al. [2014], Wang et al. [2013]. A few approaches have also been developed for the related, but more challenging problem of identifying somatic variations in matched RNA-Seq data Sheng et al. [2016].

Methods to rank and filter candidate sites are often developed using one of two approaches. Early development following the introduction of a new assay are often ad-hoc and may include the use of hard filters or other heuristic(s). These methods are widely recognized as being sub-optimal, but are nevertheless widely applied to new assays. We believe that these methods are used in these instances because: (1) they do not require a strong knowledge of probabilistic models, (2) they are easy to implement in software and (3) domain experts can look at results and contribute suggestions to improve the next iteration of the approach and (4) they enable ranking interesting candidates in the early days of an assay to demonstrate its biological interest. At these stages, it is more important to identify a few strong candidates than to obtain optimal sensitivity and specificity across a wide range of datasets.

In contrast, probabilistic methods are developed when an assay becomes popular and more data is produced that requires more sensitive or specific ranking and filtering tools. Developing probabilistic methods relies on a model of the source of errors in an assay. Developing this model for a new assay can be a slow process, but once introduced, probabilistic methods frequently outperform ad-hoc approaches by a wide margin.

In this manuscript, we describe a third approach to the ranking and filtering of genomic sites. We aimed for an approach that (1) would be fast to develop and implement for a new assay (2) would provide domain experts with the opportunity to contribute to the development and refinement of the approach, (3) would yield state of the art probabilistic models, (4) can be applied to a wide range of assays and experimental designs.

We initially developed this approach to call somatic variation in matched RNA-Seq samples. RNA-Seq is a high-throughput sequencing assay that measures gene expression Cloonan et al. [2008], but whose data can also be used to identify variation in DNA [Sheng et al. [2016] (in the parts of the genes that are expressed at sufficient level to be detectable in a given sample). A few methods have been developed to call somatic variations in RNA-Seq data, including Sheng et al. [2016] and this task is generally considered more challenging than calling somatic variations in exome or whole genome data. RNA-Seq characteristics that make the assay more challenging are (1) base coverage is unequal across the genome and driven by the expression of the genes encoded at these bases. (2) Splicing complicates genotype calling by introducing many locations in the genome (exon-exon junctions of expressed genes) where aligners may mis-align reads to the reference. (3) RNA editing is a set of biological mechanisms that modifies the sequence of RNA and can be regulated differently in different cell types. Edited bases may therefore appear as mutations in RNA when DNA is not mutated.

The main contribution of this manuscript is to present a new paradigm to develop approaches to rank and filter genomic sites. In this paradigm, we use semi-simulated datasets to train a probabilistic model. We evaluate the performance of models trained in this way for calling mutations in RNA-Seq and exome data, discuss the portability of the model from one assay to the other, and describe the versatility of the paradigm by training models for an experimental design where DNA from parents of a patient is available. In contrast to existing approaches, we propose that this new method can be used to quickly adapt a probabilistic model to specific experimental and analysis protocols.

## RESULTS

### A new paradigm to develop probabilistic models

The key idea of this study is presented in Figure 1 where we describe how we train probabilistic models using semi-simulated datasets. This approach does not rely on pre-existing training sets. Instead, we create semi-simulated datasets by planting signal in real datasets and then train probabilisitic models to recover the signal from the noise found in these datasets. We chose deep learning to implement the probabilistic models of this approach because deep neural networks (technically, networks are called “deep” if they have three or more layers) can approximate arbitrary functions and have produced state of the art performance on a wide range of machine learning problems Angermueller et al. [2016].

**Figure 1.**
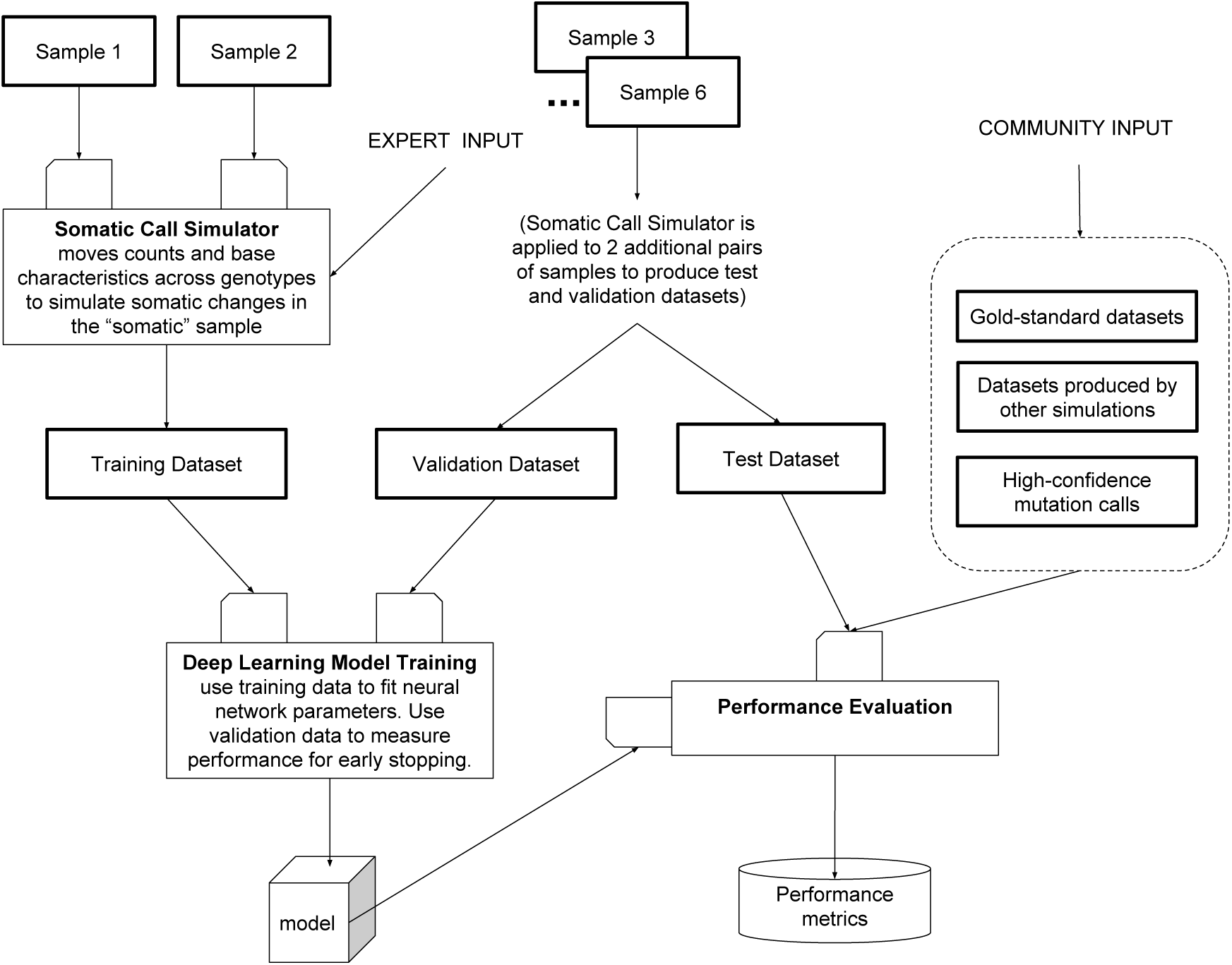
Overview of the approach. A minimum of two samples are necessary to produce a semi-simulated dataset for training. These samples are chosen from data measured from the same individual such that genotypes should match at the majority of sites measured. One sample is arbitrarily assigned the germline role, and the second sample is assigned the somatic role. These samples are provided to the somatic call simulator (whose design can be informed by expert input), which will produce mutated examples using non-mutated examples provided in the input samples (see Methods). Mutations are always added in the sample labeled “somatic”. This process yields a training dataset. The same process is repeated with independent pairs of input samples (typically from distinct subjects) to yield a validation and a test set. The training set is used to train a feedforward neural network until performance measured on a small part of the validation set starts to decrease (early stopping). Performance of the fit model can be estimated on the validation set as well as on other benchmark datasets contributed by the community.

### Need for large training sets

Training deep learning neural networks requires large training datasets with several orders of magnitude more examples than parameters in the model. Training state-of-the-art deep learning models often requires millions of training examples. However, most problems of interest in Bioinformatics have less than a fraction of a percent of the training size requirements.

### Semi-simulated training sets

To circumvent this problem, we train deep learning models with semi-simulated datasets. Semi-simulated datasets are constructed by simulating signal into samples where no signal is otherwise expected. For instance, in this project, we use two RNA-Seq samples from different types of immune cells from the same subject. Few if any somatic mutations are expected for most sites of the genome in these cells^1^. We simulate somatic variations in one of the two samples by moving bases and associated features of the real data to another genotype (see Material and Methods for details). The resulting semi-simulated dataset should accurately reflect the properties of the real data used as input, and for instance, it is expected to capture characteristics of the library preparation and sequencing assay used to obtain the data. Such characteristics include distribution of base quality for errors and somatic variations, distribution of positions in the reads where variations are observed (i.e., called read index in this study), as well as other features used to train the models.

### Training probabilistic models

An overview of this paradigm is shown in Figure 1. Briefly, we train deep neural network models using large semi-simulated training sets. The training sets are constructed using million of mutation examples constructed by planting artificial mutations in biological or technical replicates. The replicate samples are chosen to provide measured data with variability similar to that expected in future samples where the model will be applied. For instance, to call somatic variations in RNA-Seq data, we trained models using sorted immune cells from normal controls, where pairs of samples where constructed from a choice of B, T and NK cells form the same individual. We chose to use biological replicates from different cell types because somatic variations are often called in a different tissue than the germline tissue and the replicates need to reflect differences in gene expression between samples.

### Encoding Data as Features

The samples depicted in Figure 1 consist of reads aligned to a genome. In order to effectively train a deep learning network, it is necessary to convert alignments to features and labels which are compatible with back-propagation. In this work, we converted alignment data to features using the process shown in Figure 2. Briefly, alignments were realigned around indels and lined up by genomic position with the Goby framework Campagne et al. [2013, 2016b]. This conversion yields summary data about which bases and indels are observed at a given genomic site, as well as the number of reads that support the base/indel at the site. These data were then converted to 62 features per site as illustrated in Figure 2 (see Methods for details about feature mapping).

**Figure 2.**
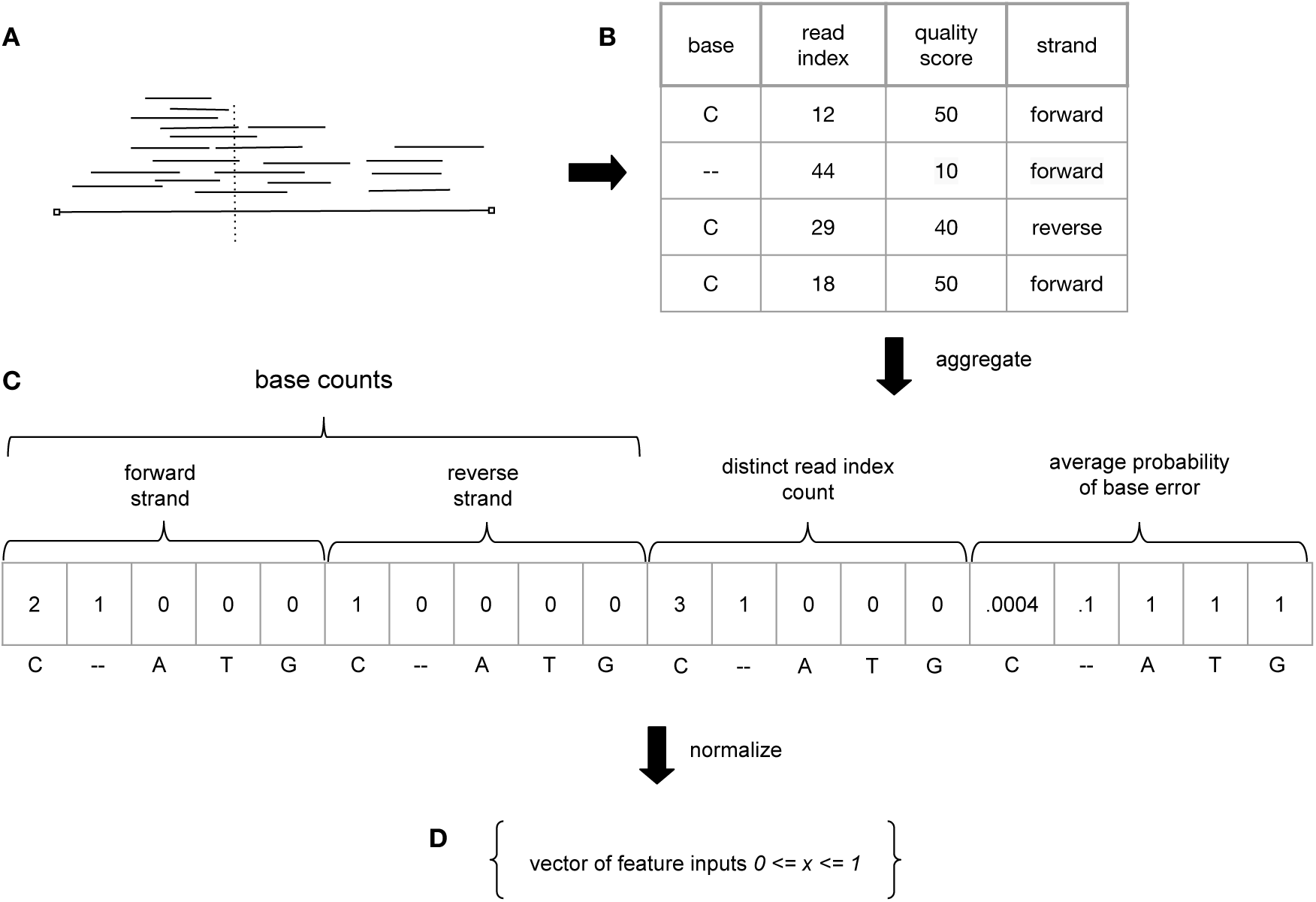
Overview of feature encoding. (A) Aligned reads are processed to obtain data about individual genomic sites (see Material and Methods for details). (B) Illustrates the data collected at each site for observed bases or indels, the genotype of the reference is available, but not shown. (C) While the number of reads at each site can vary dramatically, these data are aggregated into a fixed number of features necessary to train a neural network. The characteristics of each base observed at a site, such as, for instance, average probability of base error (derived from quality scores), number of reads and position in the read supporting the base (read index), are the result of the aggregation. (D) Aggregated features are normalized in the range 0 to 1. This figure shows the steps taken to map one sample to the feature vector (a subset of features is shown). Features derived from additional samples are concatenated.

The determination of the optimal mapping of read alignment records to features required experimentation and was performed in an iterative fashion until we found that the score on the validation set (used for early stopping) did not improve substantially. This iterative model refinement process also included manual inspection of results obtained on the validation set, to identify any unexpected predictions, and correct software bugs that can lead to them (see Figure 3). Neural network architecture and neural net hyper-parameters were also optimized during this process. The final implementation of feature mapping is modular and makes it possible to plug in or out features derived from an alignment so that different combinations can be tested and compared.

**Figure 3.**
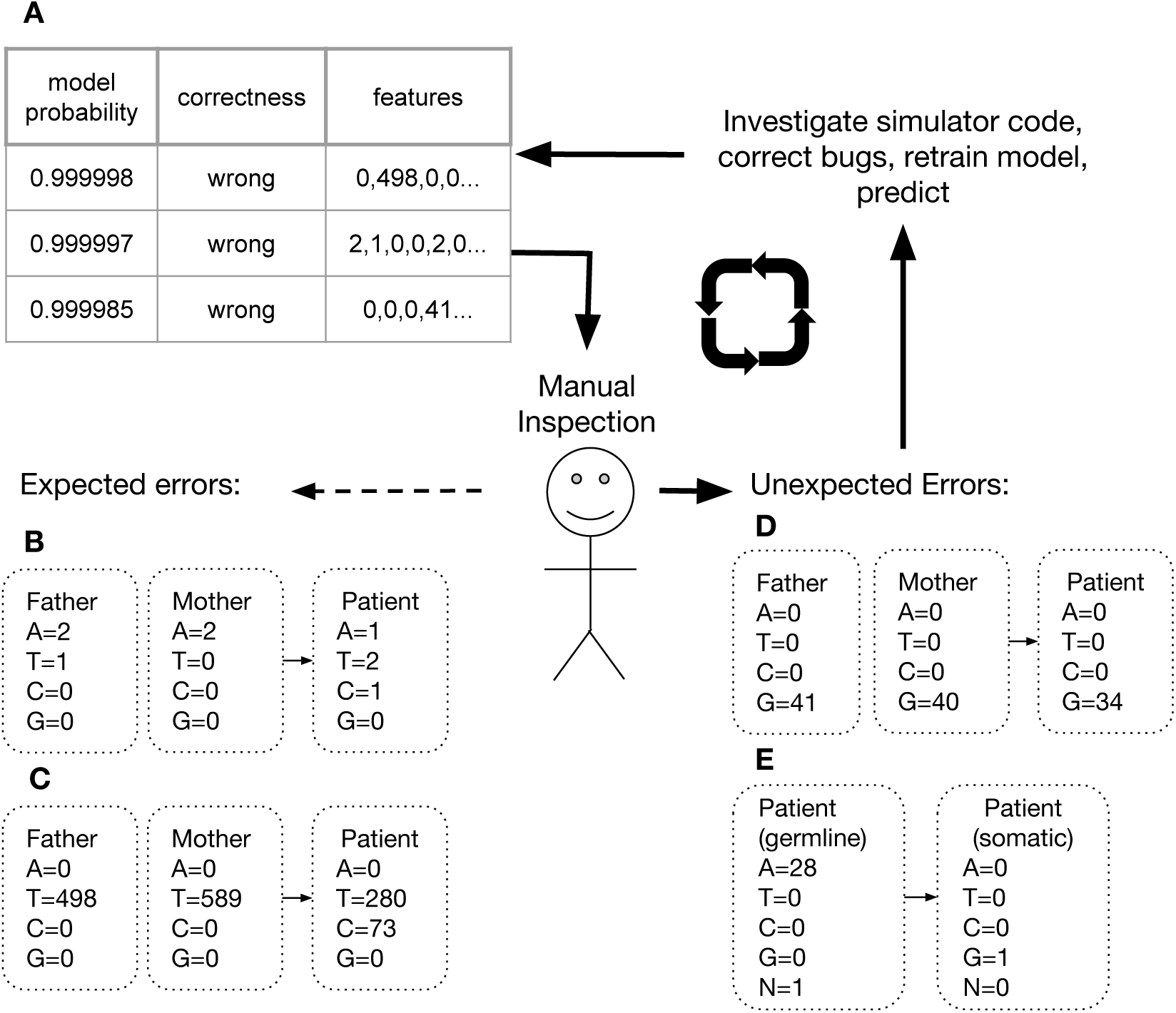
Model Refinement Cycle. (A) Example of incorrect predictions on a dataset. The table is ordered by model probability (that the site is a somatic mutation) and filtered to show only false positive predictions. Manual inspection of such a table helps identify the most extreme errors made by a model. (B) A low signal to noise ratio, as in this case, is an expected source of false positives and may indicate that a model needs further training. (C) This negative example is indistinguishable from a positive example (it looks just like the simulator planted a mutation, when this was not the case). Such errors are possible if the genotype of germline samples differ at some sites (D) This negative example was wrongly classified as a mutation. Upon further investigation of the simulator code, we identified a software bug. During the feature encoding phase, the order of bases was not consistent from sample to sample. Fixing this bug resolved such errors. (D) In a first attempt at simulation, we trained with datasets restricted to have at least 10 counts across the samples at a site. This prevented the model from learning that such sites are more difficult to predict and resulted in over-estimates of model probability. Removing filters from the construction of the training set (to include sites irrespective of their coverage) resolved this issue.

### Somatic mutation calling for an RNA-Seq assay

We applied the process shown in Figure 1 to develop models to rank and filter somatic mutations in RNA-Seq data. Briefly, we trained a model with data from two individuals consisting of pairs of control samples (B vs T, T vs B, NK vs B, B vs NK,…) where mutations were planted to create a semi-simulated dataset (see Methods for details). The model was trained with early stopping (see Methods). Performance was measured with the AUC: the area under the Receiver Operator Characteristic (ROC) curve. At the end of training, performance measured on the validation set was 0.974. A test of the model on an independent test set gave an AUC of 0.953. Table 1 summarizes the training test and validation performance of the models developed in this study.

**Table 1.** Performance of Models.

| Model | Training Set | Validation AUC | Test Set | Test AUC |
| --- | --- | --- | --- | --- |
| RNA-Seq | RNA-Seq | 0.974 | RNA-Seq | 0.953 |
| RNA-Seq | RNA-Seq | 0.974 | Pair Exome | 0.737 |
| Pair Exome | Pair Exome | 0.996 | Pair Exome | 0.994 |
| Trio Exome | Trio Exome | 0.990 | Trio Exome | 0.993 |

### Somatic mutation calling for an exome assay

An important question is whether the modeling infrastructure developed and optimized for the RNA-Seq assay can be reused to train models for a different assay.

To address this question, we trained a new model with exome data obtained with the HaloPlex exome assay (Agilent, see http://www.genomics.agilent.com/article.jsp?pageId=3081). This Haloplex assay uses enzymatic cleavage of DNA which results in very different alignment profiles than RNA-Seq protocols (which usually employ sonication to break cDNAs into fragments before adapter ligation). HaloPlex reads stack sharply at the positions where the mix of restriction enzymes cut DNA. To train the model with exome data, we used two germline samples from the same individual. These samples are expected to display germline genotypes across the entire exome. We produced semi-simulated training sets and trained an HaloPlex exome model. At the end of training, performance measured on the validation set was 0.996. A test of the model on an independent test set gave an AUC of 0.994 (see Table 1). The increase in performance of the model was expected because an exome assay is specifically optimized to produce data suitable to call genotypes, whereas RNASeq assays are optimized to measure gene expression. This result suggests that the features developed for the RNA-Seq data are transferable to a different assay when a model is retrained with a dataset for the new assay.

### Somatic mutation calling for exome and trio design

An additional question is whether the semi-simulation approach can be adapted to support different experimental designs.

To address this question, we developed models for a trio design. Assume somatic calls are needed for a patient for whom germline DNA is not readily available (because blood is suspected to contain cells with the somatic variations and contamination of other tissues by somatic cells cannot be ruled out). Assuming data from DNA from both parents of the patient are available, it should be possible to call somatic variations in the patient sample. We call this experimental design “trio somatic”, in contrast to the “pair somatic” design discussed in the previous section. We combined feature mappers developed for the pair somatic problem such that features that depend on a single sample are calculated successively for the father, mother and patient, and features that depend on two samples are calculated for father/patient and mother/patient pairs. All features were concatenated. As shown in Table 1, training this model produced similar performance to that trained for the pair exome design. This result indicates that the semi-simulation approach can train models that learn the mendelian inheritance rules necessary to call somatic variations in trio designs.

### Adaptive Models

The performance of the models developed in this study is summarized in Figure 4. Each panel shows the received operating curve (ROC) as well as a reliability diagrams Niculescu-Mizil and Caruana [2005]. The reliability diagrams compare the expected true positive rate given by the output model probability, to that observed in an independent test set (Niculescu-Mizil and Caruana [2005]). In each panel (A, C and D) where we train the model on the same experimental protocol, we find almost optimal model reliability. In Panel B, we test a model trained on RNA-Seq data on the paired exome data. The purpose of this comparison is to determine how transferable a model trained in one assay is to another assay. We find that the model remained predictive for about 40% of the exome sites, but has lower performance on the exome data. The reliability diagram also confirms that the model performs less reliably across assays. Taken together these data suggest that large performance gains can be obtained by training a model to the specific assay that it will be used for, adapting the model to the assay. While we have not measured this effect at this time, we expect that smaller performance advantages may result from adapting a model to specific data analysis protocols.

**Figure 4.**
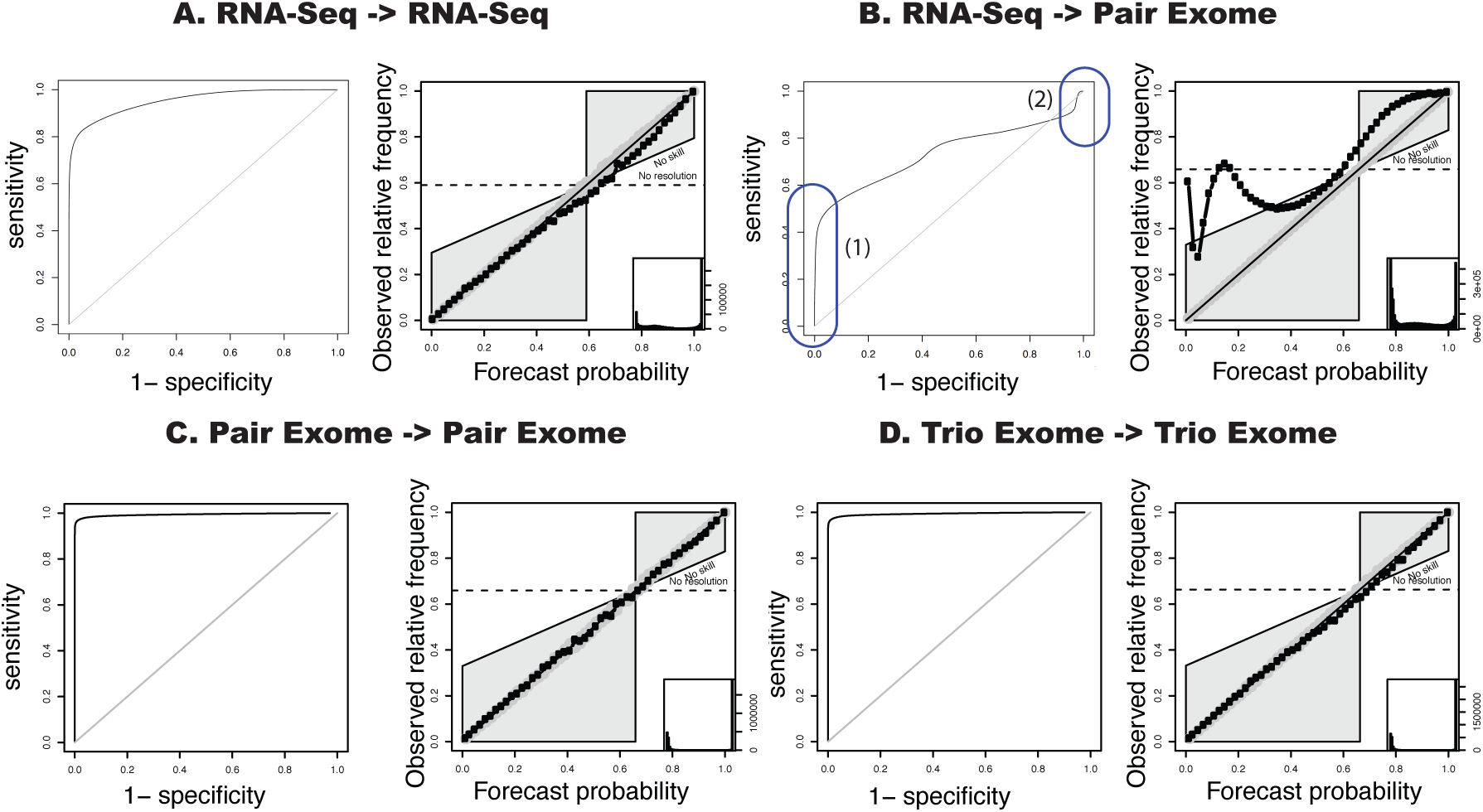
Model Performance. We characterize performance with the Receiver Operating Curves (ROC, shown on the left of each panel) and Reliability Diagrams (RC, shown on the right of each panel). (A) The model trained on RNA-Seq data performs well on RNA-Seq data and has strong reliability. (B) The same model trained on Pair Exome data predicts accurately a subset of high-confidence sites (1), with high predicted probability of mutation, but has degraded performance on other sites (2). This is confirmed by the RC, which shows sub-optimal reliability.

## Software implementation

### Using trained models

Models trained in this study have been integrated in release 3.0+ of the Goby somatic caller (distributed at http://github.com/CampagneLaboratory/goby3). Goby3 supports alignments in the Goby or BAM formats. A parameter is used to specify the path to a model to estimate probabilities of somatic variation.

### Training new models

Models for new assays can be trained by constructing a semi-simulated dataset using data obtained with the specific assay or analysis protocol. The samples used to train the model should contain no somatic variation (or a small expected number of variations when completely germline samples cannot be obtained). Detailed steps are documented with the software (see https://github.com/CampagneLaboratory/variationanalysis), but briefly, the semi-simulated dataset is produced by converting the samples to a raw dataset using the Goby SEQUENCE BASE INFORMATION output format. The file is mutated with the simulator corresponding to the experimental design (pair or trio), randomized, and split into training, validation and test sets. The training of the model uses the training and validation sets. Final model performance can be measured on the test set to check that the model generalizes.

## DISCUSSION

### Training probabilistic methods

We have presented a new approach to develop probabilistic models for calling somatic variations in high-throughput sequencing data. In contrast to previous studies which have manually designed probabilistic models, we show that it is possible to train probabilistic models using semi-simulated datasets and deep learning methods and that such models adapt to the characteristics of the data produced by specific assays and analysis protocols.

### Additional validations are needed

Additional validations will be needed to firmly establish that the models trained in this way are predictive on real datasets. Because of regulatory limitations on data sharing of genotype data, which have hampered our ability to obtain suitable validation data, we chose to distribute the software and the models to make it possible for others to conduct independent evaluations on private datasets. We hope to be able to test the models on the ICGC GoldSet Alioto et al. [2015] (request initiated with ICGC in July 2016, approved Sept 2016, we have yet to obtain access to the data files for the ICGC GoldSet which cannot be located through the ICGC portal. An email to the ICGC support mailing list to request access through the ENA, as per ICGC instructions on the ENA web site has remained unanswered as of end of Sept 2016).

### Training for new Assays and Experimental Designs

Should our approach perform competitively on real datasets, one of its key advantage will be that training can be performed for arbitrary combinations of new assays, experimental designs, and analysis protocol.

For instance, training for a new assay only requires a dataset where no somatic variations are expected (such as biological or technical replicates from the same individual). An assay-specific model can be trained following simple documented steps using the software that we have developed for this study. Training for a different analysis protocol (e.g., combination of read pre-processing and aligner) can be performed similarly.

Training for a new experimental design requires additional feature engineering, which must be implemented in new software. In our experience, developing and testing new feature mappers can be done in a couple weeks by a scientist familiar with the software when adequate computational resources are available.

### Semi-simulated datasets

Many previous studies have taken advantage of machine learning approaches to train probabilistic methods using annotated real datasets. This work differs in that we create the training sets using real datasets where no difference exists, and plant signal artificially in the real data background. If no datasets can be found with no expected difference, training could be also be performed with a dataset where only a minority of the sites are mutated. This would result in under-estimates of the probability of mutation at a site, but this bias should be small if the proportion of mutated sites in the sample is low with respect to the number of planted mutations introduced in the sample.

### Of the expected mutation rate

We showed that our approach has strong reliability when applied on the same type of assay as that used in the training set. The model outputs the probability that a site is mutated, given the prior distribution of mutated sites in the training set and the reliability diagrams indicate that the forecast probability produced by the model is close to optimal (i.e., lying very close to the diagonal on the reliability diagram). When the proportion of mutated sites in a dataset differs from that used in the training set, as is often the case in practice, it will be necessary to apply Bayes Theorem to adjust for the difference in priors. Doing so will require an estimate of the rate of mutation in the dataset where predictions must be made. Importantly, such adjustments will not change rank order, and for this reason we expect that the model probability output by the software is suitable for ranking somatic mutation in new samples.

## MATERIAL AND METHODS

### Subject characteristics and recruitment

Data from four subjects was used for developing the RNA-Seq somatic caller. RNA was obtained from sorted cells for control subjects who participated to the CFS/ME study. Any minor subject(s) (*<*18 years old in New York State) who did not provide written acknowledgment of parental permission to participate in the study where excluded. The study was reviewed and approved by the Weill Cornell Medical College Institutional Review Board (protocol 1302013563. “Immune cell gene expression and predictive models in CFS”), and patients and controls gave written informed consent after the study protocol was fully explained. All consented to blood draw and to the availability of the stored samples for additional bioassays and analyses.

Data from 12 subjects (three subjects for paired exome caller and nine subjects for trio design: three subjects and their parents) was used to develop the trio somatic caller. The study involving the exome and trio subjects was approved by the Comite´ de Protection des Personnes (CPP), Ile de France 5, (05/12/2012).

### RNA-Seq alignments

RNA-Seq reads were aligned to the 1000 Genome Project human reference sequence (corresponding to hg19) using STARR Dobin et al. integrated with GobyWeb Dorff et al. [2013].

### Exome and Trio alignments

Exome and Trio reads were aligned to the 1000 Genome Project human reference sequence as previously described Mesnard et al. [2016].

### Training, Validation and Test Datasets

The RNA-seq training, validation and test sets were created by serializing alignments to the .sbi/.sbip format with the Goby framework. The .sbi/.sbip format is a protocol buffer format developed with methods described in Campagne et al. [2013], which serializes information about aligned bases for one or more samples. The record of information in the .sbi/.sbip format is a single genomic site. For the somatic models, each record stored information about a sequenced position in two different samples (germline or somatic), or three samples (mother, father, and somatic). The protobuf schema for the .sbi/.sbip format is distributed in the Goby 3.0 repository (https://github.com/CampagneLaboratory/goby3/goby-distribution/protobuf/BaseInformationRecords.proto) Campagne et al. [2016b]. The Goby 3.0 discover-sequence-variation mode was used to serialize Goby alignments to training, validation and test datasets using the SEQUENCE BASE INFORMATION format (as described at https://github.com/CampagneLaboratory/variationanalysis).

### Mutation Simulator

The simulator is responsible for constructing millions of both positive and negative examples to train the neural network, using data records about individual genomic sites. Data records are produced by Goby and contain most of the sequencing data collected from the two samples aligned at a given genomic site (single position). The construction of mutated examples proceeds by scanning data records one at a time. First, the simulator determines if a record follows the count distribution expected of a diploid genome (canonical sites). Following this determination, only the canonical sites are mutated. The non-canonical sites are written as is with the label non-mutated. The criteria to identify canonical sites are as follows:

- the data indicate that at most two alleles are observed in either sample. This determination is made when the counts from more than 2 genotypes, with genotypes ordered in decreasing count order, have to be summed to reach 90% of all base counts for the site.
- the alleles identified by the 2 genotype rule match across samples at the site.

In this study, 18% of RNA-seq and 6% of Haloplex records were found to be non-canonical. Canonical sites are used by the simulator to create two mutated versions of the data record. The one unmutated and two mutated versions are added to the training set. Making a mutated version of a record consisted of a few steps. We used a simple heuristic to determine the original genotype, and picked a random “source” base from the two alleles in the genotype. We generated a frequency of mutation, *f*, from the uniform distribution between 0 and 0.5 (heterozygote site) or 1 (homozygote site), and chose a random “target” genotype from any of the genotypes not marked as source. We then subtracted a proportion *f* of bases from the source genotype, and added them to the target genotype. Each base retained the same features it was associated with in the source base, namely its read index location, quality score, and forward/reverse read direction.

### RNA-Seq Datasets

We used B, NK, and T-cell samples from two control subjects (CFS/ME study). For each position, a different record represented a different permutation of two cell-types for a subject, so we had 12 records for each position. We introduced mutated records with the mutation simulator, as described previously, and then randomized the order of the records within the training set.

The validation and test sets were created in the same way, using only NK and B cells data from one subject for each set.

**Table 2.** Dataset Characteristics.

|  | # Training Examples | # Validation Examples | # Test examples | # Features |
| --- | --- | --- | --- | --- |
| **RNA-seq** | 8159893 | 721012 | 666090 | 62 |
| **Pair Exome** | 2776689 | 3648105 | 1092657 | 62 |
| **Trio Exome** | 4487139 | 3907871 | 1576158 | 90 |

### Pair Exome Datasets

For pair exome datasets, we only created records of positions for one sample pairing (DNA extracted from the subject’s blood and skin). The training, validation, and test sets each drew data from an independent subject. Mutated records were introduced in training, validation and test sets. The records of each set were shuffled with the Randomizer2 tool provided in Torracinta and Campagne [2016].

### Trio Exome Datasets

For trio exome datasets, we created records of positions for one sample trio (DNA from father’s blood, mother’s blood, and subject’s blood). The training, validation, and test sets drew data from independent trios. Mutated records were introduced in training, validation and test sets. The records of each set were shuffled.

## Neural Network Architecture

### Feature Mappers

Feature mappers convert alignments about one or more samples into a fixed set of features suitable for training with neural networks. Regardless of the number of reads at a genomic position, mappers needed to produce a fixed-length output so that these outputs could be concatenated consistently into a fixed-length input vector. At each position, a mappers were used to generate the number of reads supporting each genotype (counts), the number of distinct locations in the read that support the genotype (distinct read indices), the average error probability of a base being an error (derived from base quality scores at positions that do not match the reference), and the difference in genotype proportion between the first sample (always germline) and the second sample (possibly containing a somatic mutation). For every given position, mappers which produced features about just one sample were reapplied to each sample in the assay with the results concatenated. The last mapper, which generated a comparative feature between two samples, was applied once for pair design (germline/somatic), and twice for trio design (father/child,mother/child). Mappers are implemented in the variationanalysis project available at GitHub https://github.com/CampagneLaboratory/variationanalysis.

### Model Architecture

Models were developed with the DeepLearning4J (DL4J) framework (http://deeplearning4j.org/), version 0.5.0. DL4J was selected because it is a Java framework and the models it produces can be integrated with the Goby framework more easily than frameworks in other languages. Models were formulated as 5 fully connected layers with RELU activation and a fully connected layer with soft-max activation. The first layer contained 5 times the number of input features. Inner layers contained 0.65 the number of neurons in the preceding layer. The exact model architecture used is encoded in the class called org.campagnelab.dl.varanalysis.learning.architecture.SixDense LayersNarrower2 distributed in the variationanalysis project Torracinta and Campagne [2016].

### Training Procedure

Stochastic gradient descent optimization was used, with an ADAGRAD optimizer (called updater in the DL4J framework). The model trained on mini-batches of 600 examples. Regularization was not used.

Finding that it did not introduce training instability, the learning rate was set to the starting value of 1, with SCORE decay (the learning rate was decreased when the loss on the validation set stopped decreasing).

An early stopping condition of 10 epochs interrupted training early if no performance improvement was observed in the validation set after 10 training epochs. Models were trained with early stopping by measuring performance on 100,000 examples from the validation set.

### Performance measurements

AUC measures were calculated with the org.campagnelab.dl.varanalysis.stats.AreaUnderTheROCCurve class, provided in the in the variationanalysis project Torracinta and Campagne [2016]. This class implements the naive *O* (*n*^2^) calculation of the area under the ROC curve and directly calculates the probability of correctly classifying an observation. This class was adapted from the AUC calculator implemented in the BDVal project, which was validated in the MAQC-II project Consortium et al. [2010].

Received Operating Curves were produced with the AUC R package, integrated in MetaR Campagne et al. [2016a]. Reliability diagrams were constructed with the SpecsVerification R package, integrated in MetaR Campagne et al. [2016a].

## AUTHOR CONTRIBUTIONS

RT and FC wrote the variation analysis programs as well as parts of Goby3 necessary to create datasets. LM provided exome and trio data and developed detailed protocols for blood collection and processing in the CFS/ME study. SL helped obtain the four control samples of the CFS/ME project used to train and validate models in this study. RS developed detailed protocols for B cell, T cell and NK cell sorting in the CFS/ME study and supervised blood extraction, cell sorting and RNA extraction. MH and FC supervised the human subject components of the CFS/ME study. FC designed the computational approaches.

## ACKNOWLEDGMENTS

We thank Manuele Simi for technical assistance with the code needed for this project.

## Footnotes

1 Exceptions are the sites that code for B and T cell receptor sites, which undergo somatic mutations in these cell types, but are still a minority among all the sites of a genome.

## REFERENCES

Tyler S Alioto, Ivo Buchhalter, Sophia Derdak, Barbara Hutter, Matthew D Eldridge, Eivind Hovig, Lawrence E Heisler, Timothy A Beck, Jared T Simpson, Laurie Tonon, et al. A comprehensive assessment of somatic mutation detection in cancer using whole-genome sequencing. Nature communications, 6, 2015.

Christof Angermueller, Tanel Pärnamaa, Leopold Parts, and Stegle Oliver. Deep Learning for Computational Biology. Molecular Systems Biology, (12): 878, 2016.

Fabien Campagne, Kevin C. Dorff, Nyasha Chambwe, James T. Robinson, and Jill P. Mesirov. Compression of Structured High-Throughput Sequencing Data. PLoS ONE, 8(11):e79871, nov 2013. ISSN 1932-6203. doi: 10.1371/journal.pone.0079871. URL http://dx.plos.org/10.1371/journal.pone.0079871.

Fabien Campagne, William ER Digan, and Manuele Simi. Metar: simple, high-level languages for data analysis with the r ecosystem. bioRxiv, page 030254, 2016a.

Fabien Campagne, Remi Torracinta, and Manuele Simi. Goby 3.0.0 software release, 2016b. URL https://doi.org/10.5281/zenodo.159024.

Nicole Cloonan, Alistair R R Forrest, Gabriel Kolle, Brooke B a Gardiner, Geoffrey J Faulkner, Mellissa K Brown, Darrin F Taylor, Anita L Steptoe, Shivangi Wani, Graeme Bethel, Alan J Robertson, Andrew C Perkins, Stephen J Bruce, Clarence C Lee, Swati S Ranade, Heather E Peckham, Jonathan M Manning, Kevin J McKernan, and Sean M Grimmond. Stem cell transcriptome profiling via massive-scale mRNA sequencing. Nature methods, 5(7):613–9, 2008. ISSN 1548-7105. doi: 10.1038/nmeth.1223. URL http://www.ncbi.nlm.nih.gov/pubmed/18516046.

MAQC Consortium et al. The microarray quality control (maqc)-ii study of common practices for the development and validation of microarray-based predictive models. Nature biotechnology, 28(8): 827–838, 2010.

A Dobin, C A Davis, F Schlesinger, J Drenkow, C Zaleski, S Jha, P Batut, M Chaisson, and T R Gingeras. doi: bts635[pii]10.1093/bioinformatics/bts635.

Kevin C. Dorff, Nyasha Chambwe, Zachary Zeno, Manuele Simi, Rita Shaknovich, and Fabien Campagne. GobyWeb: Simplified Management and Analysis of Gene Expression and DNA Methylation Sequencing Data. PLoS ONE, 8(7):e69666, 2013. ISSN 1932-6203. doi: 10.1371/journal. pone.0069666. URL http://www.pubmedcentral.nih.gov/articlerender.fcgi?artid=3720652{\&}tool=pmcentrez{\&}rendertype=abstract.

Laurent Mesnard, Thangamani Muthukumar, Maren Burbach, and Carol Li. Exome Sequencing and Prediction of Long-Term Kidney Allograft Function. pages 1–15, 2016. doi: 10.1371/journal.pcbi. 1005088.

Alexandru Niculescu-Mizil and Rich Caruana. Predicting good probabilities with supervised learning. In Proceedings of the 22nd international conference on Machine learning, pages 625–632. ACM, 2005.

Stephan Pabinger, Andreas Dander, Maria Fischer, Rene Snajder, Michael Sperk, Mirjana Efremova, Birgit Krabichler, Michael R Speicher, Johannes Zschocke, and Zlatko Trajanoski. A survey of tools for variant analysis of next-generation genome sequencing data. Briefings in bioinformatics, 15(2):256–78, mar 2014. ISSN 1477-4054. doi: 10.1093/bib/bbs086. URL http://www.pubmedcentral.nih.gov/articlerender.fcgi?artid=3956068{\&}tool=pmcentrez{\&}rendertype=abstract.

Quanhu Sheng, Shilin Zhao, Chung I. Li, Yu Shyr, and Yan Guo. Practicability of detecting somatic point mutation from RNA high throughput sequencing data. Genomics, 107(5):163–169, 2016. ISSN 10898646. doi: 10.1016/j.ygeno.2016.03.006.

Remi Torracinta and Fabien Campagne. Variationanalysis 1.0.2 software release, October 2016. URL https://doi.org/10.5281/zenodo.159203.

Qingguo Wang, Peilin Jia, Fei Li, Haiquan Chen, Hongbin Ji, Donald Hucks, Kimberly Brown Dahlman, William Pao, and Zhongming Zhao. Detecting somatic point mutations in cancer genome sequencing data: a comparison of mutation callers. Genome medicine, 5(10):91, 2013. ISSN 1756-994X. doi: 10.1186/gm495. URL http://www.pubmedcentral.nih.gov/articlerender.fcgi?artid=3971343{\&}tool=pmcentrez{\&}rendertype=abstract$\backslash$nhttp://www.ncbi.nlm.nih.gov/pubmed/24112718$\backslash$nhttp://www.pubmedcentral.nih.gov/articlerender.fcgi?artid=PMC3971343.

